# FISHnet: Detecting chromatin domains in single-cell sequential Oligopaints imaging data

**DOI:** 10.1101/2024.06.18.599627

**Authors:** Rohan Patel, Kenneth Pham, Harshini Chandrashekar, Jennifer E. Phillips-Cremins

## Abstract

Sequential Oligopaints DNA FISH is an imaging technique that measures higher-order genome folding at single-allele resolution via multiplexed, probe-based tracing. Currently there is a paucity of algorithms to identify 3D genome features in sequential Oligopaints data. Here, we present FISHnet, a graph theory method based on optimization of network modularity to detect chromatin domains and boundaries in pairwise distance matrices. FISHnet uncovers cell type-specific domain-like folding patterns on single alleles, thus enabling future studies aiming to elucidate the role for single-cell folding variation on genome function.

## Main

A decade of technology development leading to chromosome-conformation-capture sequencing assays^1–4^ have revealed that the mammalian genome is folded into A/B compartments, topologically associated domains (TADs), subTADs, and loops^5–8^. Using bulk sequencing assays and perturbative experiments, leading models assert that TADs/subTADs and their boundaries regulate gene expression by restricting the physical search space of enhancers for their distal target genes and preventing ectopic enhancer-promoter interactions (reviewed in^9^). Boundary disruption has been linked to gene expression dysregulation in multiple models of human disease^10–12^. Moreover, boundaries created by cohesin-mediated loop extrusion stalling at high density arrays of CTCF binding sites have been functionally linked to the placement of replication initiation zones in early S-phase^13^. A major question emerging from bulk Hi-C-based studies is whether domain-like folding patterns can be detected in single cells. To further elucidate the genome’s structure-function relationship, there is a need for technologies to quantitatively detect TAD/subTAD folding patterns at single cell resolution.

Multiplexed sequential DNA FISH Oligopaint imaging technologies have recently enabled single-allele imaging of genome folding at kilobase resolution across Megabase (Mb)-sized sections of the genome^14–18^. Oligopaints imaging experiments use tiled probes and sequential imaging steps to generate dense matrices of pairwise distances between genomic loci, thus allowing the direct visualization of genome folding in single alleles and across a population of single cells (**Fig. 1a**)^15^. Advances in computational methods to detect domain-like patterns in single-allele imaging data have been slowed by key challenges. First, the signal to noise ratio is low within pairwise distance matrices which makes sensitive and specific domain detection while minimizing false positives particularly challenging. Second, the signal in distance matrices represents a small dynamic range of possible measured nanometer distances between genomic loci on the order of 0-1 micrometers. By contrast, signal in Hi-C sequencing data can range from 0 up to 1 million or more sequencing reads per valid pair of contacts. Third, Oligopaints experiments involve tiling of probes in a stepwise bin-by-bin manner and are prone to a high number of dropouts due to technical biases such as low chromatin accessibility of probes or low probe specificity due to repetitive regions of the genome. The dropouts are often interpreted by algorithms as boundaries, which increases false positive domain calls. In bulk Hi-C data, this particular challenge is overcome because a missed proximity-ligation event in one cell averages out across millions of cells. Moreover, bins with low mappability in Hi-C matrices can be computationally corrected with Knight-Ruiz matrix balancing. Overall, the challenges inherent to sequential Oligopaints data underscore the need for specialized analytical approaches tailored to the unique characteristics of single-allele resolution datasets distinct from those of bulk Hi-C technologies.

**Figure 1.**
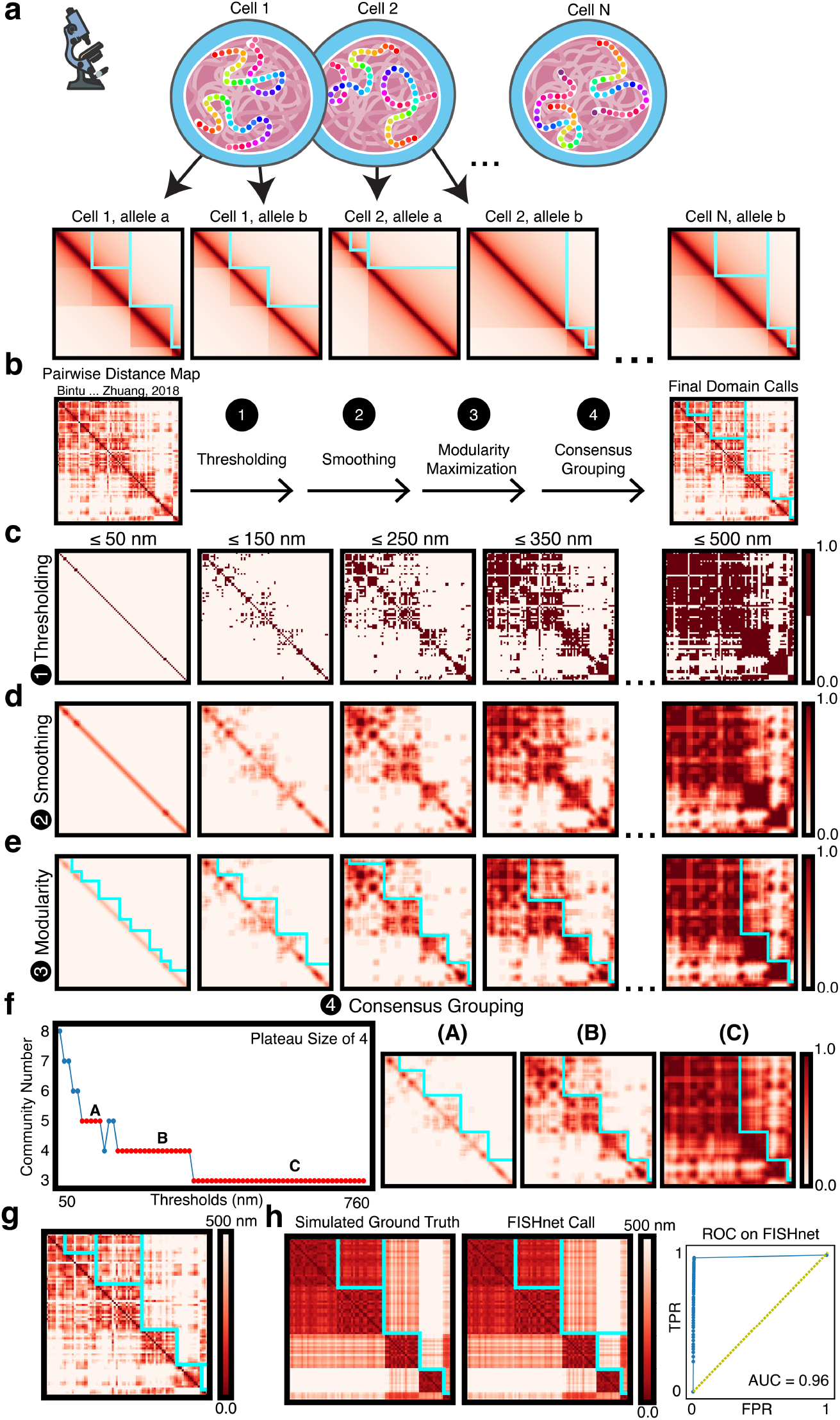
Schematic of domain-like pattern detection in single-cell sequential Oligopaints imaging data with FISHnet. **(a)** Cartoon schematic of sequential Oligopaints data and simulated pairwise distance matrices with FISHnet chromatin domain calls overlaid. **(b)** FISHnet methodological steps. The pairwise distance matrix of a 30 kb resolution trace spanning chr21:34.6Mb-37.1Mb on a single-allele from HCT116 cells^15^. **(c)** Binary matrices resulting from a sweep of nm distance thresholds on the input pairwise distance matrix. **(d)** Smoothing of binary matrices. **(e)** Domain calls per each smoothed, binarized matrix by optimizing network modularity (see Methods). **(f)** Domain calls as a function of distance threshold. Those within the same plateau were merged and shown overlaid on averaged smoothed, binarized matrices from each distance-thresholded group. **(g)** Final non-redundant domain calls overlaid on the pairwise distance matrix. **(h)** ROC curve measuring FISHnet’s performance on simulated pairwise distance matrices. Yellow dotted line showcases y=x.

To detect domain-like patterns in sequential Oligopaint data, we developed FISHnet as an open-source graph theory-based method that employs four steps (binarization with thresholding, smoothing, network modularity maximization, and consensus grouping) to identify chromatin domains (**Fig. 1b**). FISHnet begins by transforming a 2D array of nm pairwise distance measurements between genomic loci into multiple binarized matrices after thresholding (**Fig. 1c**). Thresholding is a technique to isolate entries within the pairwise distance matrix that have distance values that are less than or equal to the threshold value by converting them into a binary value. FISHnet uses multiple thresholds since different distance thresholds give rise to different domain structures present within the pairwise distance matrix allowing FISHnet to capture different degrees of domains. We select the range of distance thresholds by computing the minimum and maximum pairwise distances found within the matrix converting the array to binary values after thresholding on distance in a gradient of 10 nm steps. We discovered that small distance thresholds such as 50 or 100 nm yield binarized arrays with small and nested domain-like patterns and large distance thresholds such as 500 nm or greater yield larger and encompassing domain-like patterns (**Fig. S1**). In addition to revealing domains of different sizes, thresholding helps increase the signal-to-noise ratio which increases sensitivity and specificity of downstream computational steps.

We next smoothed the binary matrices by applying a 2x2 signal averaging window to the binarized matrices (**Fig. 1d**). Smoothing aims to convert the binary matrices into a dynamic range of 0 to 1 and represents bins with high interactions with each other with respect to the distance threshold. We apply a small window to prevent boundary shifting. Hereafter, we refer to the binarized and smoothed arrays as adjacency matrices. We hypothesized that chromatin domains can be identified in adjacency matrices through a community detection method based on the maximization of network modularity (*Q*) (**Equation (1)**):

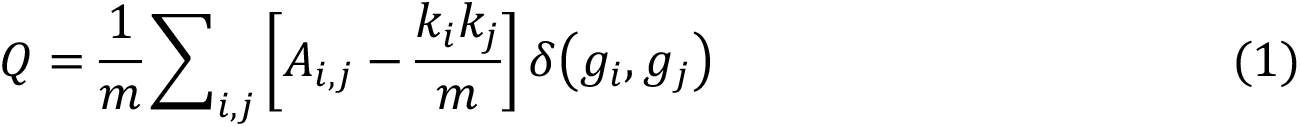

where *A* is the adjacency matrix, *A*_*i,j*_ is the edge weight representing the interaction between nodes *i* and *j*, *k_i_* is the sum of all of the edge weights for node *i,* and *m* is the sum of all non-diagonal edge weights within the network *A*. Nodes *i* and *j* are assigned to communities *g*_*i*_ and *g*_*j*_ respectively. The Kronecker delta, δ(*g*_*i*_, *g*_*j*_), ensures that only nodes that are within the same group when maximizing modularity are considered in the calculation. It is 1 if *g*_*i*_ = *g*_*j*_ and 0 otherwise.

We used the modularity equation to evaluate if two nodes are interacting, independent, or not interacting via a computed observed – expected relationship. The observed value is computed *A_ij_* as and the expected value is computed as 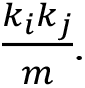. For values of modularity, a positive non-zero observed-expected calculation >0 represents two interacting nodes and signifies domains whose members are close in physical distance to each other and far from members of other domains. A zero observed-expected calculation represents the nodes that are independent from each other and thus exhibit similar intra- and inter-domain distances. A negative non-zero observed-expected calculation <0 represents two nodes that do not interact. Thus, a high modularity metric is conceptually intuitive as a metric for quantifying chromatin domains because they are regions of the genome that fold within physical proximity to members within its domain versus outside the domain^19^.

We apply a Louvain-like algorithm^20^ (see **Methods**) to the adjacency matrices to maximize modularity (**Fig. 1e**). For each adjacency matrix the Louvain-like algorithm is applied 20 times to account for stochasticity in the maximization algorithm. At each nm distance threshold, we computed a consensus set of domain calls from 20 runs of modularity maximization using the adjusted RAND score metric^21, 22^ (see **Methods**). We then plotted the number of domains (or communities) as a function of nm distance threshold. When the plot plateaus at 4 or more adjacent nm distance steps, we compute a consensus partition representing the plateau and then merge all consensus partitions as the final domain calls (**Fig. 1f-g**).

We assessed FISHnet’s accuracy in calling domain-like structures by simulating pairwise distance matrices with known domain locations (**Methods, Figure S2**). Using N=1000 simulations we applied FISHnet and computed an area under the curve (AUC) of 0.96 from the Receiver Operating Characteristic (ROC) curve (**Fig. 1h**). A common flaw in sequential Oligopaints data is pairwise distance matrices in which the majority of genomic locations have dropouts due to poor probe efficiency. We simulated pairwise distance matrices with a range of 5-50% bin dropout (**Figure S3a**). Moreover, we used linear imputation as previously published^24^ to smooth over the dropouts (**Figure S3b**). We observed that FISHnet is robust for data with up to 20% dropouts with an AUC of 0.91 (**Figure S3a**). Linear interpolation improves FISHnet’s performance on simulated data with known domains and dropouts to an AUC of 0.89 with 40% of the map missing (**Figure S3b**). Our data demonstrate that FISHnet is sensitive and specific for calling domains in single- allele Oligopaints data, even in scenarios with substantial missing data.

We next tested the extent to which FISHnet performance corresponds to boundaries present in ensemble Hi-C data. We ran FISHnet on publicly available sequential Oligopaints data from human HCT116 cells at 30 kb resolution^15^ and mouse embryonic stem (ES) cells at 25 kb resolution^18^. Given FISHnet’s low false positive rate with linear imputation on maps with up to 40% of bin dropouts, we ran FISHnet with linear imputation on all single alleles with pairwise distance maps at more than 60% coverage. Representative pairwise distance matrices from each dataset along with their respective FISHnet calls (**Fig. 2a**) demonstrate FISHnet’s ability to identify domain structures from pairwise distance matrices. Using bulk Hi-C data^25, 26^ and ensembled sequential Oligopaints data, we also demonstrate that the FISHnet boundary calls are present in the highest proportion of cells when they are also present in bulk Hi-C data (**Fig. 2b**). We further find that the architectural proteins CCCTC-binding factor (CTCF) and cohesin subunit RAD21^25, 27^ correlate with the frequency of domain calls in single-alleles (**Fig. 2b**). Cohesin and CTCF are architectural proteins that fold the genome via loop extrusion^19, 28, 29^. Members of the cohesin complex form a ring-like structure which progressively moves along DNA, extruding out the intervening DNA into a loop until the ring stalls at boundary elements bound by CTCF. The extrusion process is the mechanistic basis for loops and the manifestation of topologically associated domains (TADs), and subTADs in ensemble Hi-C data^19, 30–32^. We observe a strong enrichment for CTCF/RAD21 peaks at close proximity to single-allele boundary calls (**Figure S4**). Together, our work reveals a link among Hi-C, CTCF and RAD21 peaks, and the frequency of FISHnet single-allele boundary calls in sequential Oligopaints data. Our work supports the sensitivity and specificity of FISHnet’s domain calling capability in single-allele imaging data.

**Figure 2:**
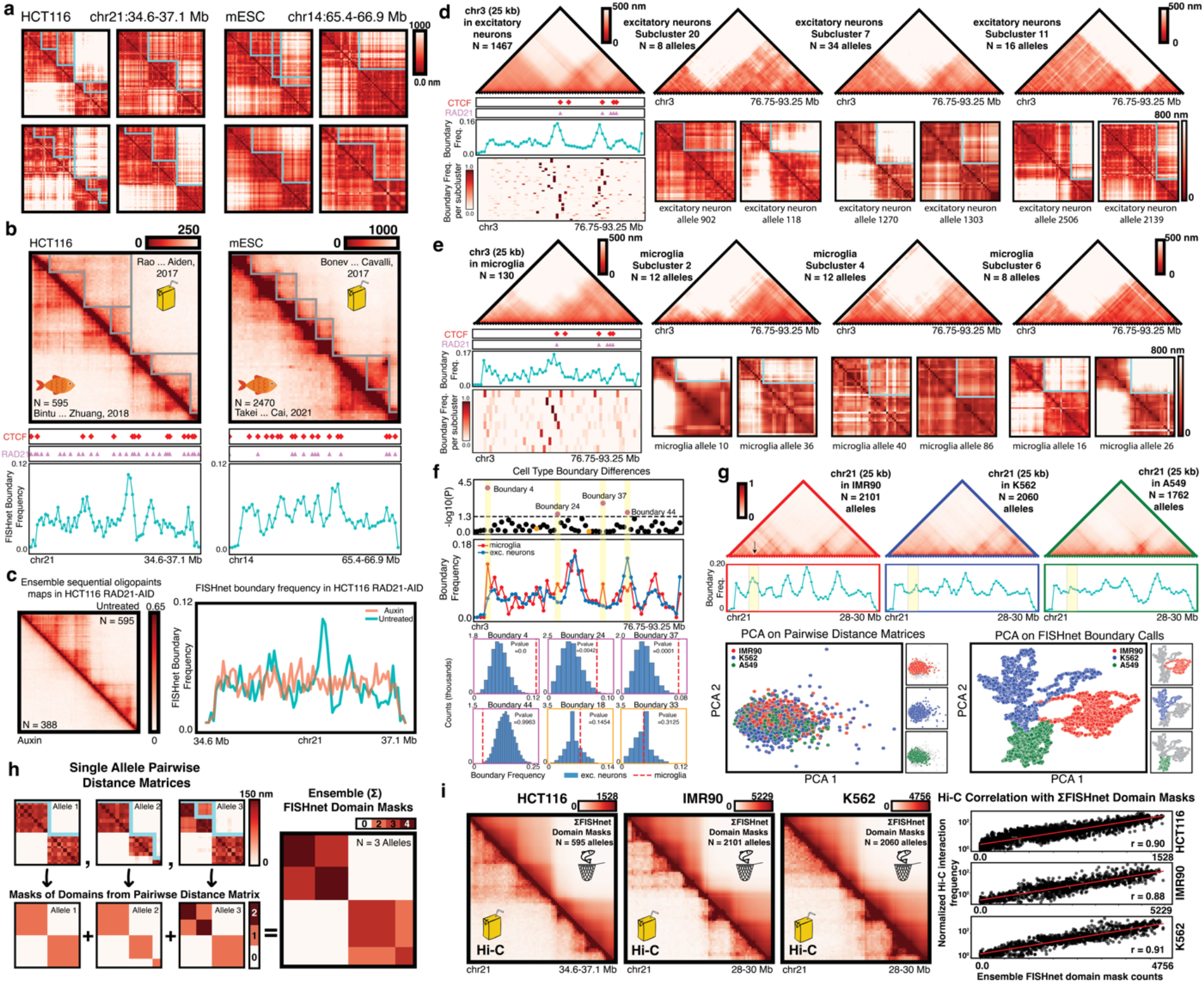
Results and applications of FISHnet. **(a)** Individual pairwise distance matrices with FISHnet calls on published sequential Oligopaints imaging data in mouse embryonic stem (mES) cell alleles and HCT116 alleles. Color bar indicates nm pairwise distances between genomic loci. Mouse ES alleles were traced at 25 kb resolution while HCT116 alleles were traced at 30 kb resolution. **(b)** Hi-C with TAD/subTAD calls using 3DNetMod^23^ in mES cells (25 kb resolution) and HCT116 cells (30 kb resolution) is provided in the top half of the heatmaps overlaid with CTCF and RAD21 peak calls. The ensemble counts matrices of sequential Oligopaints imaging data from the same cell type is provided in the bottom half of the maps (N=595 alleles, HCT116 and N=2470 alleles, mESC). Color bar indicates normalized interaction frequency (Hi-C) and number of alleles with distances less than 250 nm (DNA FISH). Blue line plots indicate the frequency of FISHnet boundary calls across N= 595 HCT116 alleles and N = 688 mESC cell alleles. **(c)** Ensemble frequency matrices of sequential Oligopaints imaging data in HCT116 RAD21-AID cells in the untreated (N = 595 alleles) and auxin-treated (N = 388 alleles) conditions at 30 kb resolution. Color bar indicates number of alleles with distances less than 250 nm normalized by the total number of alleles. Adjacent to the heatmap are line plots representing the frequency of FISHnet boundary calls in single cells from untreated and auxin-treated conditions. **(d-e)** Ensemble averaged matrices of all sequential Oligopaints imaging data in (**d**) excitatory neurons (left, N=1467 alleles) and (**e**) microglia (left, N=130 alleles) as well as subclusters of alleles with different boundary structures (three right excitatory neuron ensemble averaged maps, N=8, 34, and 16 alleles, respectively; three right microglia ensemble averaged maps, N=12, 12, and 8 alleles, respectively). Below each ensemble map are single-allele example heatmaps of sequential Oligopaints imaging data representing that particular domain pattern. Domain calls on single-allele pairwise distance matrices are of the largest nm plateau calls. Blue line plots representing the frequency of FISHnet boundary calls in single cells overlaid with CTCF and RAD21 peak calls from mouse brain tissue. Color bar indicates nm pairwise distance between genomic loci. Oligopaints data for excitatory neurons and microglia were traced at 25 kb resolution. **(f)** Pvalues for differential boundaries for every bin within excitatory neurons and microglia using the chi-squared test. Underneath is line plots representing the frequency of FISHnet boundary calls for excitatory neurons and microglia. For boundaries significant within the chi-square test a permutation test via bootstrapping is shown. Two negative permutation tests are also shown in orange. **(g)** Ensemble frequency matrices of sequential Oligopaints imaging data in IMR90 (N = 2101), K562 (N = 2060), and A549 (N = 1762) alleles. Color bar indicates number of alleles with distances less than 250 nm normalized by the total number of alleles. Blue line plots represent the frequency of FISHnet boundary calls for each cell type respectfully. Yellow vertical highlights indicate IMR90 cell type specific boundary. Principal component analysis on the three human cell lines showing the first two eigenvectors run on using pairwise distance matrices and FISHnet boundary calls. **(h)** Cartoon schematic of method for implementing an ensemble FISHnet domain mask. **(i)** Ensemble FISHnet domain mask arrays (top) compared to ensemble Hi-C data (bottom) for HCT116, IMR90, and K562 alleles. Color bar indicates normalized interaction frequency (Hi-C) and ensembled FISHnet domain mask counts (DNA FISH). Scatterplots of the normalized Hi-C interaction frequency and the ensemble FISHnet domain mask counts for each cell type are shown. Red line indicates the linear line of best fit, and Pearson correlation values are indicated for each scatterplot.

We sought to understand if FISHnet performance is robust across data from a perturbative model system. We ran FISHnet on sequential Oligopaints data generated in human HCT116 cells engineered with an auxin inducible degron system tagging RAD21^15^, mouse brain tissue^33^, and human IMR90, K562, and A549 cells^15^. Upon auxin treatment to remove RAD21, we observed that RAD21 depletion results in an equal probability of FISHnet boundary calls across the locus^15^ (**Fig. 2c**). These results are consistent with the finding of lost TAD/subTAD boundaries genome wide upon RAD21 disruption^25^.

To test if FISHnet could detect differential boundary positions among cell types within tissue, we used published Oligopaints imaging data from mouse brain tissue^33^. We ran FISHnet on sequential Oligopaints data from both excitatory neurons and microglia and plotted the frequency of boundary calls overlaid with CTCF and cohesin peak calls and ensembled imaging matrices (**Fig. 2d-e**). We first ran FISHnet on sequential Oligopaints pairwise distance matrices from N=1467 excitatory neurons (**Fig. 2d top**) and N=130 microglia (**Fig. 2e top**). We dropped maps with less than 60% coverage for excitatory neurons from further analysis and applied linear imputation before running FISHnet. We used all N=130 microglia pairwise distance maps and applied linear imputation before running FISHnet. Using FISHnet’s domain calling output, we classified Oligopaints matrices into those with boundary patterns exhibiting strong correlation with each other (see **Methods**). We found N=49 clusters with unique folding patterns for excitatory neurons (**Fig. 2d**, **Fig. S5**) and N=8 unique folding clusters for microglia (**Fig. 2e, Fig. S6**). We observed a strong correlation between the visual location of boundaries and the frequency of FISHnet domain calls in single cells in each cluster. Together, these data indicate that FISHnet has ability to resolve cell type differences and single-allele differences in boundary locations in sequential Oligopaints imaging data.

We next explored single-cell variation in genome folding between two cell types at any given boundary location. For neurons and microglia, we formulated a statistical test in which we tested the association of a FISHnet boundary call with cell type (see **Methods**). For every bin in mouse brain tissue Oligopaints data, we used a chi-squared test to compute the probability of obtaining single-allele boundary frequencies equal to or more extreme than FISHnet values if the null hypothesis is true that the boundary occurs in a similar proportion of alleles in both cell type A and cell type B. The chi-squared test effectively identified boundaries that are significantly more likely to be present in a higher proportion of single alleles in either microglia or excitatory neurons (**Fig. 2f**, **top purple points**).

We next developed an empirical permutation test that identifies microglia-specific boundaries after accounting for single-allele variation. We formulated a test statistic representing the proportion of alleles with a specific FISHnet boundary (see **Methods**). We computed a one-tailed, right-tailed empirical Pvalue by comparing the test statistic from N=130 single microglia alleles to a null distribution representing 10,000 draws of N=130 alleles from excitatory neuron Oligopaints data. Applying our permutation test, we confirmed that the boundaries identified as cell type-specific with the chi-squared test were also present in a significantly higher proportion of alleles in microglia than expected by chance from the excitatory neuron null distribution (**Fig. 2f**, **bottom**). Visually apparent neuron-specific boundaries did not result in low Pvalues as expected given the one-tailed, right-tailed test. We also chose two random boundaries not significant from the chi-square test and confirmed that they were not significant with the permutation test (**Fig. 2f**, **top orange points**). Our analyses indicate that FISHnet has capability to identify genomic locations of statistically significant cell type-specific boundaries.

We sought to understand if Oligopaints pairwise distance matrices or FISHnet boundaries could be used to discern cell types. We ran FISHnet on published sequential Oligopaints imaging data from N=2101, N=2060, and N=1762 single alleles from IMR90, K562, and A549 cells, respectively^15^. Plotting the ensemble single-allele data demonstrated that the visual location where specific boundaries are present in IMR90 cells and absent in K562 and A549 cells correlates with a high FISHnet boundary single-allele frequency only in IMR90 (**Fig. 2g**, top left). Principal component analysis using the pairwise Oligopaints imaging distance matrices could not distinguish cell types. However, principal component analysis using the single-allele FISHnet boundary calls alone could readily separate IMR90 from K562 and A549 (**Fig. 2g**, bottom). Altogether, our data reinforce FISHnet’s ability to identify cell type-specific boundaries across a range of cell lines and its ability to distinguish cell type-relevant genome folding features more effectively than the raw data.

FISHnet is unique from many 3D genome methods in that it calls not only boundaries but also full chromatin domains at single-allele resolution. We tested the extent to which FISHnet domain calls correspond to TADs/subTADs present in ensemble Hi-C data (**Fig. 2h-i**). We transformed the single-allele domain call outputs from FISHnet from HCT116, K562, and IMR90 cell lines into an integer mask that represents and distinguishes domain structures within a pairwise distance matrices (see **Methods**). To create an ensemble of FISHnet domain masks, we summed all masked pairwise distance matrices (**Fig. 2h**, **bottom**). Each pixel in the ensemble FISHnet domain mask represents the number of times that it was present in a domain (**Fig. 2h**, **right**). We observed strong correlation between the ensemble FISHnet domain mask counts and ensemble Hi-C data in the same locus and cell type (Pearson’s correlation coefficient 0.90, 0.88, and 0.91 for HCT116, IMR90, and K562, respectively) (**Fig. 2i**). Together, these data highlight FISHnet’s ability to identify both boundaries and domains in a cell type-specific manner that recapitulate ensemble Hi-C data, which opens up future opportunities to understand how single-allele domain calls can yield ensemble measurements.

FISHnet is a graph-theory based method that employs modularity maximization for the sensitive and specific detection of domain-like genome folding patterns in sequential DNA FISH Oligopaints imaging data. FISHnet quantifies the frequency of boundaries in single-cells and can evaluate statistically significant cell type specific domains within pairwise distance matrices from single-alleles.

## Acknowledgments

We thank members of the Cremins lab for helpful discussions. FISHnet art used in Fig. 2i is created by Laymik from the Noun Project. K. Pham is a recipient of a F30 Ruth L. Kirschstein National Research Service Award Individual Predoctoral Fellowship.

## Author contributions

Conceptualization: RP, JEPC Methodology: RP, JEPC Investigation: RP, KP, HC, JEPC

Visualization: RP, KP, HC, JEPC Funding acquisition: JEPC Project administration: JEPC Supervision: JEPC

Writing & Editing: RP, KP, JEPC

## Declaration of interests

N/A

**FISHNet code is provided at bitbucket and zenodo links below for private reviewer access during the review process. The code links and usage instructions will be made publicly available upon publication.**

## Methods

### Hi-C binning, bias correction, and 3DNetMod

We downloaded Hi-C matrices for HCT116^25^ and mESC^26^ from GEO (**Table S1**) and for K562 and IMR90 cell lines from the 4DN portal (**Table S1**). We binned the Hi-C matrix for HCT116, K562, and IMR90 at 30 kb resolution and for mESCs at 25kb resolution. All Hi-C matrices underwent Knight-Ruiz balancing for bias correction. We called TADs/subTADs using 3DNetMod as we have previously reported with minor modifications^34–40^.

### CTCF and RAD21 peak calling

We used CTCF and RAD21 ChIP-seq peakcalls from publicly available datasets in mouse brain tissue^41^, mESCs^42^, and HCT116^36^ cell lines. For mouse brain tissue, we downloaded previously published ChIP-seq fastq files for CTCF (GSM5253974) and RAD21 (GSM5253990) and their corresponding inputs (GSM5253982, GSM5253994). We aligned CTCF and RAD21 ChIP-seq reads to the mouse (mm10) genome using Bowtie^43, 44^ (version 0.12.7). We removed reads with multiple possible alignments using -m2 flag and removed optical and PCR duplicates using samtools (version 1.2). We downsampled RAD21 pulldown and input libraries to 22 million reads and identified peaks using MACS2^45^ (version 2.1.2) broad peak calling parameters --broad -- broad-cutoff 1E-4 at p-value 1E-4. For CTCF ChIP-seq, we applied MACS2 narrow peak calling parameters at p-value 1E-4. For HCT116 cell lines, we used published peak calls for CTCF and RAD21^36^. For mES cell lines, we used published mouse mm10 peak calls from (GSE125129) for CTCF and RAD21 ChIP-seq.

### Community detection via modularity maximization (Louvain-like algorithm)

We have previously described the details of the methods in other manuscript^23^, and they are again described in detail here to ensure reproducibility. Mathematical terms and equations should be expected to be similar due to reproducibility of the methodological steps. To partition binarized and smoothed pairwise distance matrices (also referred to as adjacency matrices) into communities, we utilized a Louvain-like, locally greedy algorithm to maximize modularity. We first calculated the modularity matrix, M, according to **Equation 2**:

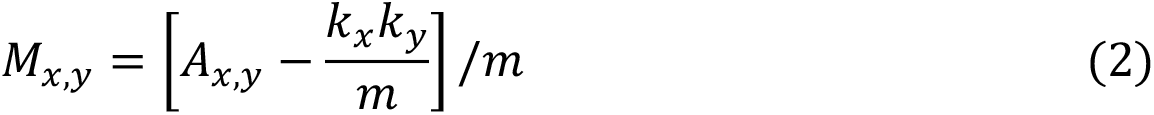

where M is the modularity matrix with size C x C and C is the number of communities. *M*_*xy*_ is the normalized interaction between community x and y, *k*_*x*_ is the sum of all edge weights for community x, and *m* is the sum of all interactions in the adjacency matrix A excluding the diagonal. In Equation (2) we see that *A*_*x,y*_/*m* is the normalized edge strength connecting communities *x* and *y* and 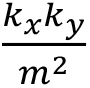 is the expected normalized edge strength at communities *x* and *y*. We computed modularity, *Q*, on the modularity matrix, *M*, with**Equation 3**:

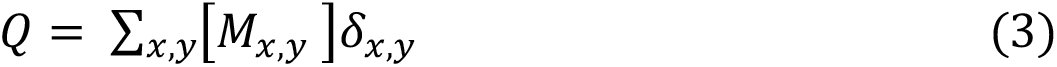

where δ_*x,y*_ ensures that only edge weights within communities are added to the summation by being 1 if *x* and *y* are assigned to the same community (*x* = *y*) and 0 otherwise (*x* ≠ *y*).

The Louvain-like algorithm works by using an iterative approach to rapidly converge on a local modularity maximum without comprehensively examining the entire search space. For each iteration, *itr*, individual nodes are given the opportunity to move into a new community placement that yields a locally maximal gain in Δ*Q* according to **Equation 4**:

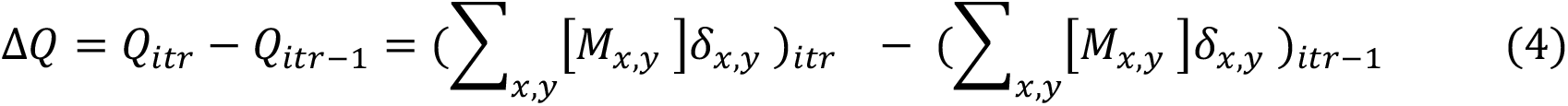

For the first iteration of the Louvain-like algorithm, every individual node is its own community. In this *itr = 0* circumstance, the indices for communities x and y correspond to the indices for nodes *i* and *j* and the modularity matrix, *M,* is the same size as the adjacency matrix *A*. Before the end of the *itr,* each node has the chance to merge to form a new community. In *itr* = 1, modularity is recalculated and if the gain in modularity (Δ*Q*) is greater than or equal to 10^−10,^, the algorithm advances to the next iteration and adjusts the dimensions of the modularity matrix, M, to C × C (where C represents the number of communities computed in the previous iteration, and (*M*_*x,y*_)*_itr_* is the sum of all previous (*M*_*x,y*_)*_itr-1_* constituent edges that are merged into (*M*_*x,y*_)*_itr_*). The algorithm terminates when no further single community merge leads to an improvement in modularity, i.e. the gain in modularity (Δ*Q*) is less than 10^*+,^. To counteract the potential convergence on local maxima inherent in modularity-maximization algorithms, the Louvain-like algorithm underwent 20 iterations. To ensure a diverse exploration of the landscape and avoid bias towards specific solutions, we randomly selected a node with a distinct seed value for each of the 20 partitions. Finally, a consensus partition is selected by taking the partition that is most similar to the 19 other partitions through the use of adjusted RAND score (see below).

### Consensus Partition through the adjusted RAND score

We have previously described the details of the methods in our other manuscript^23^, and they are again described in detail here to ensure reproducibility. Mathematical terms and equations should be expected to be similar due to reproducibility of the methodological steps. FISHnet uses the adjusted RAND score to find a consensus partition when running the Louvain-like algorithm 20 times to prevent fixation on a local maximum as well as when finding a consensus partition within a plateau sweep. The consensus partition is a partition that is most similar to the other partitions within its group. We find the consensus partition by employing a similarity metric called the adjusted RAND (aRAND) index, derived from the sklearn toolbox sklearn.metrics.adjusted_rand_score. The aRAND index yields unity for perfectly matched partitions and tends towards 0 or slightly negative values for partitions that are no more similar than expected by random chance. We computed aRAND for all pairs of partitions within the set. We chose the partition with the highest average aRAND as the consensus partition because it most resembled all other partitions.

### Creation of Ensemble counts matrix

To compute an ensemble counts matrix as seen in the bottom half of the heatmap in **Fig. 2b** we start by binarizing/thresholding all of the pairwise distance matrices within a dataset by the value of less than or equal to 250 nm. We then sum all of the binarized maps which yields the ensemble counts matrix.

### Creation of Ensemble frequency matrix

To compute an ensembled frequency matrix as seen in **Fig. 2c** and **Fig. 2g** we start by computing the ensemble counts matrix. Then to normalize for dropouts and number of pairwise distance matrices, we divide each entry in the ensemble counts matrix by the number of non-Nan elements in that entry for all of the pairwise distance matrices. This yields the ensemble frequency matrix.

### Creation of Ensemble average matrix

To compute an ensemble average matrix as seen in **Fig. 2d/e** we start by adding all of the pairwise distance matrices within a dataset and dividing the value by the total number of pairwise distance matrices in the dataset.

### Simulating Pairwise Distance Matrices

To simulate pairwise distance matrices, we follow three key steps: (1) a random walk, (2) boundary creation, and (3) random shuffling of points within domains (**Fig. S2**). The random walk is created by first starting at point (0,0,0). The next point is generated as a random point within a sphere with radius of 100 nm centered at the previous point. Steps are continued until the pairwise distance matrix size matches the intended number of bins. To create boundaries, we take all points upstream of the boundary and calculate their centroid. We then adjust all of the coordinates towards the upstream centroid. A similar procedure is done for points downstream of the boundary. Nested boundaries are then created by repeating this process within an already created domain. For this study we randomized the number of boundaries depending on the length of the simulation. We used simulations with length sizes of 42, 60, or 83. For maps with 42 or 60 bins there is a chance for 1-2 boundaries being created. For maps with 83 bins, 1-3 boundaries could be created. We also randomly shuffle coordinates within a domain to remove any artifactual domain-like patterns created by the random walk (**Fig. S2**).

### Receiver Operator Curve (ROC) creation

We computed a receiver operator curve (ROC) for FISHnet boundary calls from simulated data to test the sensitivity and specificity of the algorithm. First, we ranked each FISHnet boundary call by the number of times it appeared within its plateau grouping and created a confusion matrix for all of the pairwise distance matrices in a simulated dataset. In the confusion matrices, we counted FISHnet boundaries within 1 bin of a ground truth boundary as a true positive (TP). We classified a FISHnet boundary as a false positive (FP) when FISHnet called a boundary with no ground truth boundary within 1 bin. Then, we count a false negative (FN) when FISHnet failed to call a boundary at a ground truth boundary. Finally, true negatives (TN) represent when neither FISHnet called a boundary nor a ground truth boundary exists. We calculated the false positive rate calculated as FP/(FP+TN) and the true positive rate as TP/(TP+FN). We plotted the ROC curve with true positive rate on the x-axis and the false positive rate on the y-axis.

### FISHnet excitatory neuron and microglia subcluster classification

To identify subclusters within the excitatory neuron and microglia population as shown in **Fig. 2d**- **e** we used FISHnet calls associated with the largest nm plateau calls. We constructed an *N×M* matrix for both excitatory neurons and microglia, where *N* represents a genomic bin and *M* corresponds to a single allele. Here, *N* is the total number of genomic bins within the pairwise distance matrix, and *M* is the total number of alleles. For each allele in the *N×M* matrix, a value of 1 indicates the presence of a boundary at that bin Ni in that allele Mj, while a value of 0 indicates its absence. Then, we computed the correlation matrix to determine the Pearson correlation coefficients between each pair of alleles, transforming the *N×M* matrix into an *M×M* correlation matrix. Using Python’s ‘networkx package’, we applied the ‘greedy_modularity_maximization’ module to identify clusters within the correlation matrix. We used the ‘networkx.algorithms.community.greedy_modularity_communities’ command where *G* is the *M×M* correlation matrix thresholded at either 0.75 or 0.5 for excitatory neurons or microglia, respectively. We retained clusters comprising more than three pairwise distance matrices and discarded the rest.

### Chi-Square test for identifying cell type-specific boundaries

To identify boundaries that are significantly different between two cell types, we conducted a chi-square test of independence for FISHnet calls from each genomic locus using Oligopaints data from microglia and excitatory neurons (**Fig. 2f**). For a given boundary we constructed a 2 by 2 contingency table in which we classified and tallied each microglia and neuron single allele as either having or not having the boundary as measured by FISHnet. We used the python package Scipy to run scipy.stats.chi2_contingency(contingency_table) to test the null hypothesis that the boundary occurs in similar proportions of alleles in both cell type A and cell type B. The alternative hypothesis is that the boundary does not occur in similar proportions across alleles in both cell type A and cell type B.

### Permutation test for identifying cell type-specific boundaries

We formulated a permutation test to identify microglia-specific boundaries (cell type A-specific boundaries) as shown in **Fig 2f**, **bottom**. We selected a test statistic of the proportion of alleles with a FISHnet called boundary at a particular genomic locus. Using the microglia and neuron data, we elected to run a one-tailed, right-tailed test for the null hypothesis that the proportion of microglia and excitatory neuron alleles with a boundary is unchanged. The alternative hypothesis is that the proportion of microglia alleles with a boundary is higher than the proportion of neuron alleles with a boundary. We compute the test statistic using the microglia data and compare it to a null distribution formulated by randomly sampling 10,000x, without replacement, N=130 alleles from the excitatory neuron single allele Oligopaints data. We compute a one-tailed, right-tailed empirical Pvalue as the area under the curve of the null distribution to the right of the microglia test statistic.

### Pairwise distance matrix PCA for IMR90, K562, and A549 alleles

We performed principal component analysis on the pairwise distance matrices in **Fig. 2g** using N = 2101, 2060, and 1762 pairwise distance matrices for IMR90, K562, and A549, respectively. We removed pairwise distance matrices with missing values resulting in N = 2082, 2032, and 1700 alleles for IMR90, K562, and A549, respectively. We then extracted the distances from each pair of genomic loci and created a matrix where each row stores the distances between genomic loci. Thus, in our matrix sized *N × M*, rows *N* represent individual alleles and columns *M* contained the distances between two genomic loci for each allele. We used sklearn.preprocessing.scale to zero mean scale the matrix, calculated the covariance matrix, and computed the eigenvectors and eigenvalues using numpy.linalg.eigh. We ranked the eigenvalues and selected the eigenvectors corresponding to the two largest eigenvalues and created a reduced eigenvector matrix. Finally, we reduced the scaled matrix by taking the dot product with the transposed scaled matrix and the reduced eigenvector matrix and plotted the first two principal components.

### FISHnet PCA for IMR90, K562, and A549 alleles

We performed principal component analysis on a masked domain representation array from FISHnet domain calls in **Fig. 2g** also using N = 2101, 2060, and 1762 pairwise distance matrices for IMR90, K562, and A549, respectively. For this analysis, we used FISHnet domain calls with the smallest nm plateau to focus on nested domains because they are known to be cell type-specific. We created a *N × M* matrix as described in the “FISHnet excitatory neuron and microglia subcluster classification” section for IMR90, K562, and A549 single alleles. Next, we smoothed each column in the *N × M* matrix, where *M* stores the FISHnet boundary calls for a given allele, with a window size of 150. Otherwise, we lose contextual information such that statistical methods would give boundaries that are close to each other equal weight as boundaries farther apart. We then concatenated each of the *N × M* matrices together creating a matrix that contains the smoothed boundary calls for each cell type for all of the alleles present in the IMR90, K562, and A549 datasets. We used sklearn.preprocessing.scale to zero mean scale the smoothed matrix, calculated the covariance, and computed the eigenvectors and eigenvalues using numpy.linalg.eigh. We ranked the eigenvalues and selected the eigenvectors corresponding to the two largest eigenvalues and created a reduced eigenvector matrix. Finally, we reduced the scaled matrix by taking the dot product with the transposed scaled matrix and the reduced eigenvector matrix and plotted the first two principal components.

### Ensemble FISHnet domain mask creation

We first transformed each individual pairwise distance matrix by creating an integer mask based on each map’s FISHnet domain calls (**Fig. 2h**, **left)**. The resultant mask transforms the nm pairwise distances into an integer encoding how many domains FISHnet finds each pair of genomic loci (**Fig. 2h**, **bottom left)**. If a pixel is not in any FISHnet domain, its value in the mask is 0. If a pixel is in only one FISHnet domain, its value in the mask is 1. In cases where a pixel is in two FISHnet domains, such as one nested within a another larger one, its value is 2. To create an ensemble FISHnet domain mask, we summed individual masks, resulting in a matrix in which each pixel represents the count of domains FISHnet found each pair of genomic loci in (**Fig. 2h**, **right**).

### FISHnet Settings

For all sequential Oligopaint datasets, we ran FISHnet run with a plateau sweep of 4, size exclusion of 3, merge of 3, and smoothing window size of 2x2. We used size exclusion to require communities to be greater than 3. The merge parameter dictated the averaging of boundaries within 3 bins of each other.

