## Supplemental Methods and Figures for "FISHnet: Detecting chromatin domains in single-cell sequential Oligopaints imaging data"

Supplementary Information

Rohan Patel[1,2,3], Kenneth Pham[1,2,3], Harshini Chandrashekar[1,2,3], Jennifer E. Phillips-Cremins[1,2,3*]

1 Department of Bioengineering, University of Pennsylvania, Philadelphia, PA
2 Epigenetics Institute, Perelman School of Medicine, University of Pennsylvania
3 Department of Genetics, Perelman School of Medicine, University of Pennsylvania
* Corresponding author

**Supplementary Table 1.** Table of publicly available datasets used within this study.

| Dataset | Description | Link |
|---|---|---|
| Multiplexed sequential OligoPaints imaging data | HCT116-RAD21-AID auxin induced | https://github.com/BogdanBintu/ChromatinImaging/tree/master/Data |
| Multiplexed sequential OligoPaints imaging data | HCT116-RAD21-AID control | https://github.com/BogdanBintu/ChromatinImaging/tree/master/Data |
| Multiplexed sequential OligoPaints imaging data | mESC | https://zenodo.org/records/3735329 |
| Multiplexed sequential OligoPaints imaging data | Mouse Brain Tissue | https://zenodo.org/records/4708112 |
| Multiplexed sequential OligoPaints imaging data | K562 | https://github.com/BogdanBintu/ChromatinImaging/tree/master/Data |
| Multiplexed sequential OligoPaints imaging data | IMR90 | https://github.com/BogdanBintu/ChromatinImaging/tree/master/Data |
| Multiplexed sequential OligoPaints imaging data | A549 | https://github.com/BogdanBintu/ChromatinImaging/tree/master/Data |
| In situ Hi-C | HCT116-RAD21-AID control | https://data.4dnucleome.org/experiment-set-replicates/4DNES3QAGOZZ/ |
| Hi-C | mESC | https://www.ncbi.nlm.nih.gov/geo/query/acc.cgi?acc=GSE96107 |
| In situ Hi-C | IMR90 | https://data.4dnucleome.org/experiment-set-replicates/4DNES1ZEJNRU/ |
| In situ Hi-C | K562 | https://data.4dnucleome.org/experiment-set-replicates/4DNESI7DEJTM/ |
| CTCF Chip-seq | HCT116 | https://www.nature.com/articles/s41586-022-04803-0 supplementary table 13 |
| CTCF Chip-seq | mESC | https://www.ncbi.nlm.nih.gov/geo/query/acc.cgi?acc=GSE125129 |

| | | |
|---|---|---|
| CTCF Chip-seq | Mouse Brain Tissue | https://www.ncbi.nlm.nih.gov/geo/query/acc.cgi?acc=GSE35140 |
| RAD21 Chip-seq | HCT116 | https://www.nature.com/articles/s41586-022-04803-0 supplementary table 13 |
| RAD21 Chip-seq | mESC | https://www.ncbi.nlm.nih.gov/geo/query/acc.cgi?acc=GSE125129 |
| RAD21 Chip-seq | Mouse Brain Tissue | https://www.ncbi.nlm.nih.gov/geo/query/acc.cgi?acc=GSE35140 |

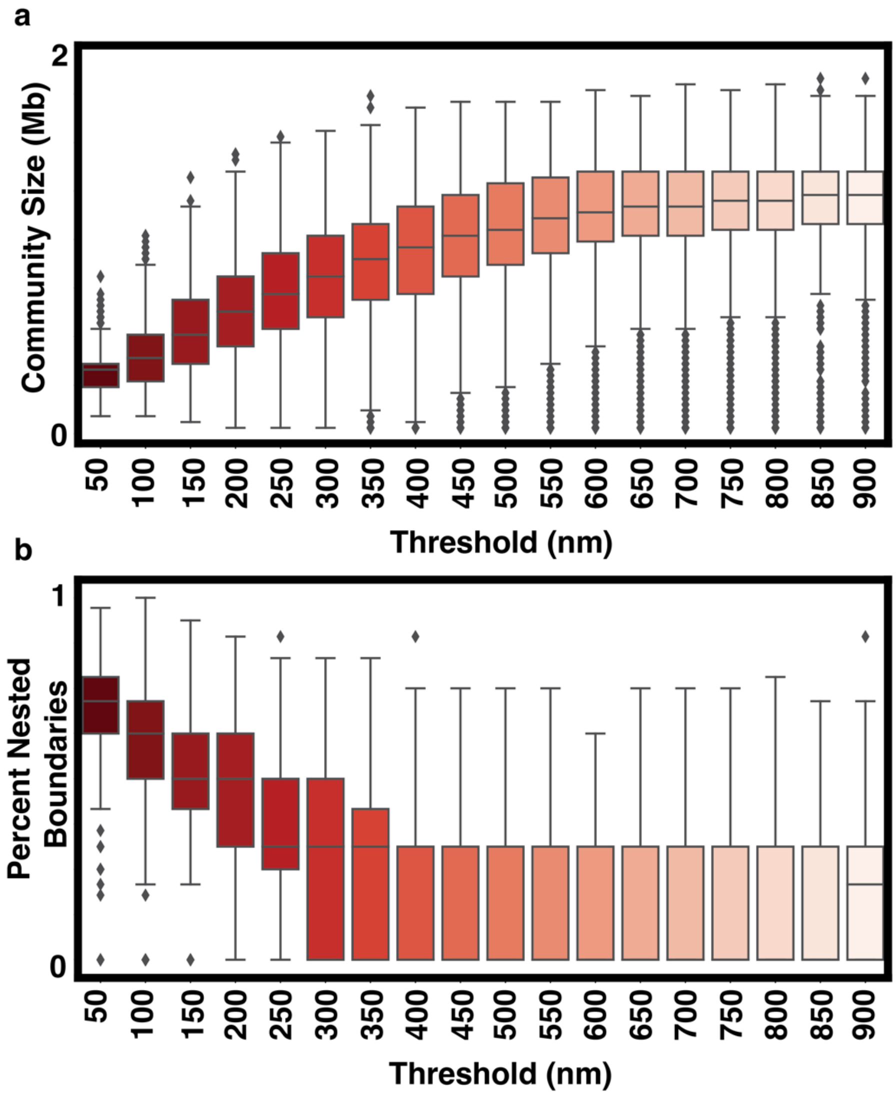


**Supplementary Figure 1. Use of nanometer distance threshold as a resolution parameter. (a)** Boxplot of domain call sizes at different nm distance threshold values for a chromatin tracing data in HCT116 cells (Chr21: 34.6–37.1 Mb at 30 kb resolution) (Bintu, Mateo … Boettiger, Zhuang, 2018). N = 595 pairwise distance maps were used. **(b)** Boxplot of percent nested boundaries at different distance thresholds from the same dataset used in (a).

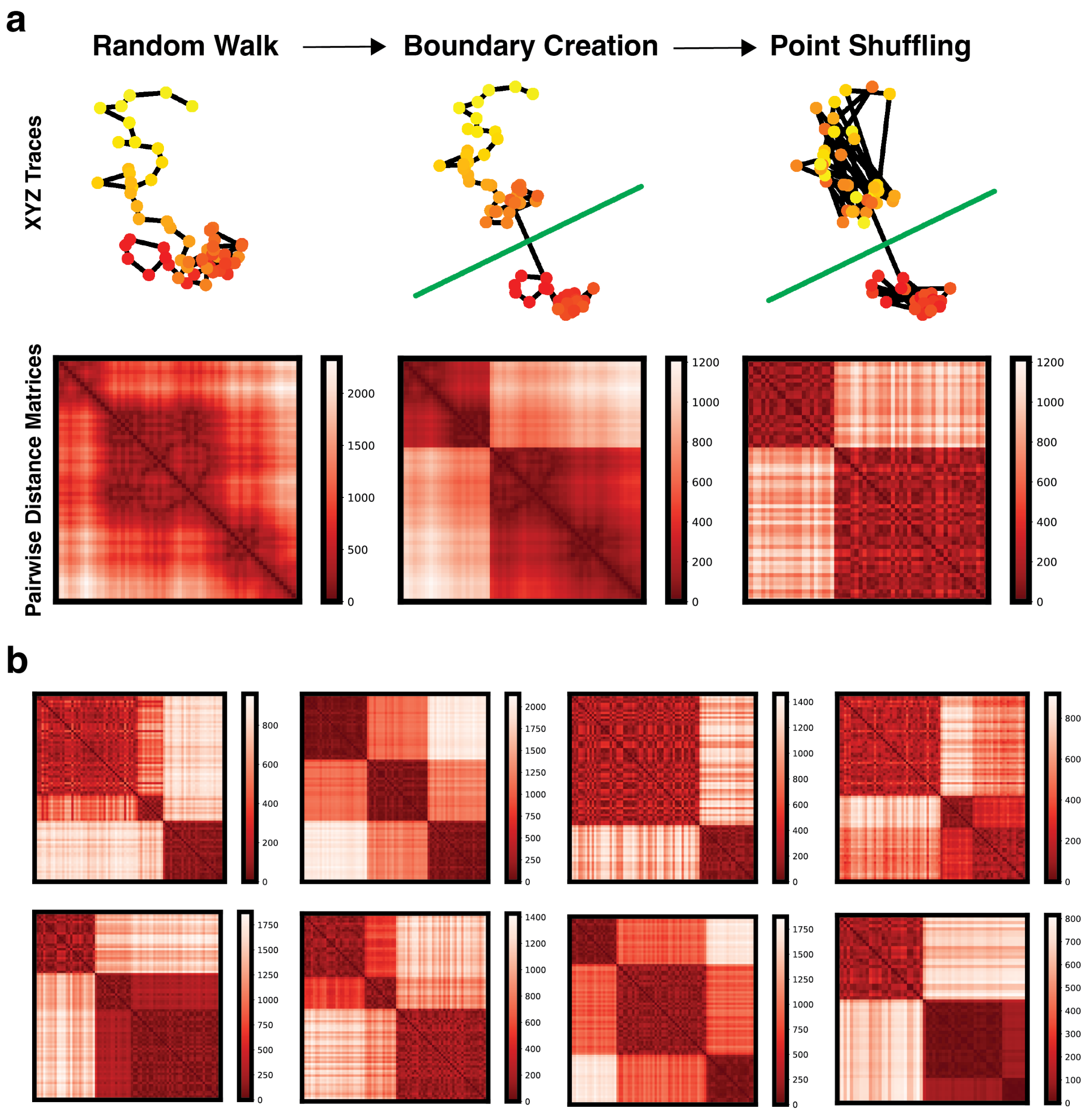


**Supplementary Figure 2. Simulated single allele oligopaints pairwise distance matrices. (a)** Schematic of steps to generate simulated oligopaints imaging maps. We simulate a random walk with N discrete points and create boundaries by forcing random points within a cluster to move closer to their respective centroid. Green lines indicate the location where the code suggests creating a boundary. Coordinates within domains are randomized to prevent unwanted domains from forming due to the random walk procedure. **(b)** Representative examples of simulated pairwise distance matrices at different complexities.

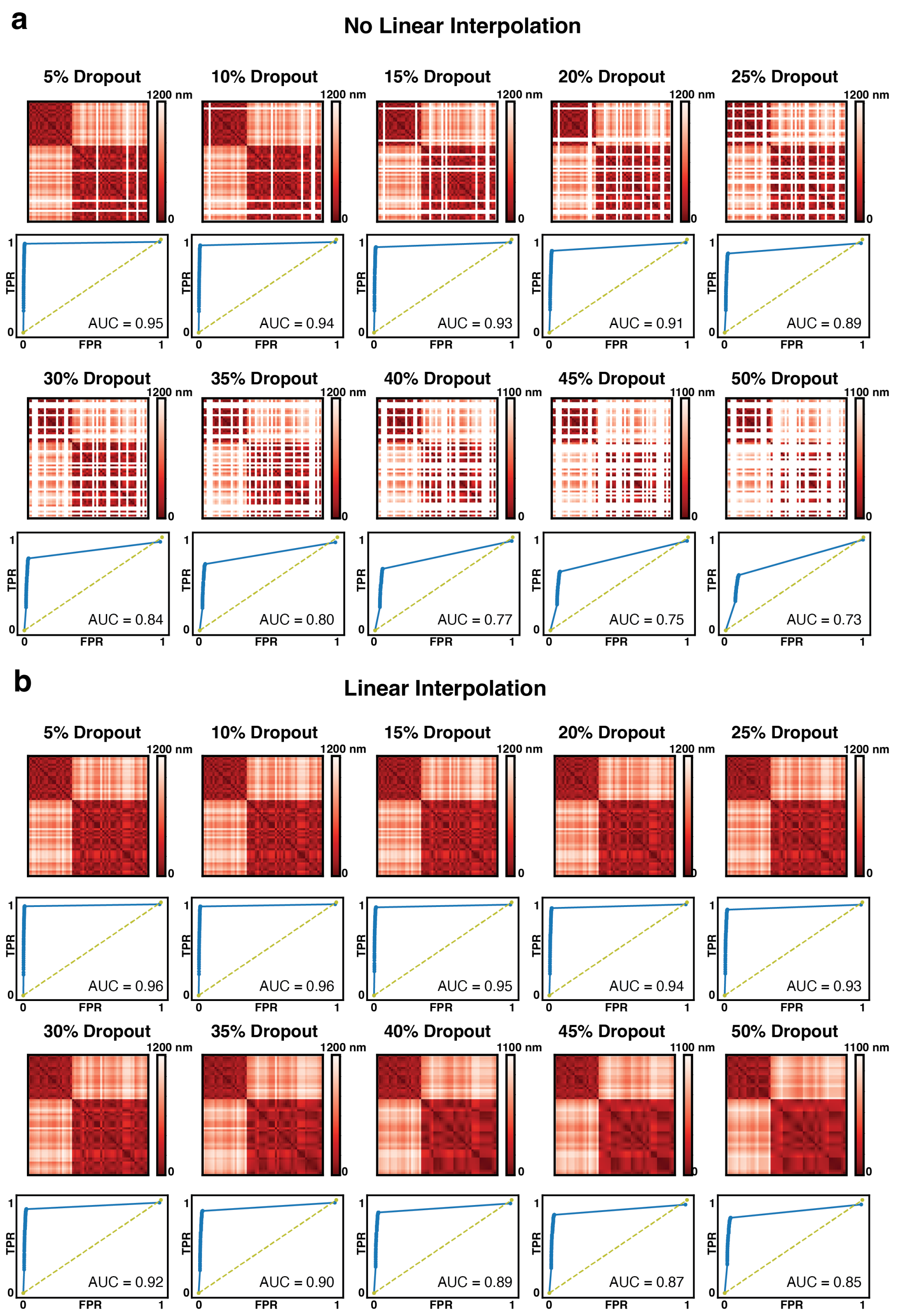


**Supplementary Figure 3. Assessment of FISHnet's sensitivity and specificity across a range of bin dropout and with and without linear interpolation. (a)** Visualization of dropouts with ROC curves for FISHnet's performance without linear interpolation. **(b)** Visualization of dropouts with ROC curves for FISHnet's performance with linear interpolation.

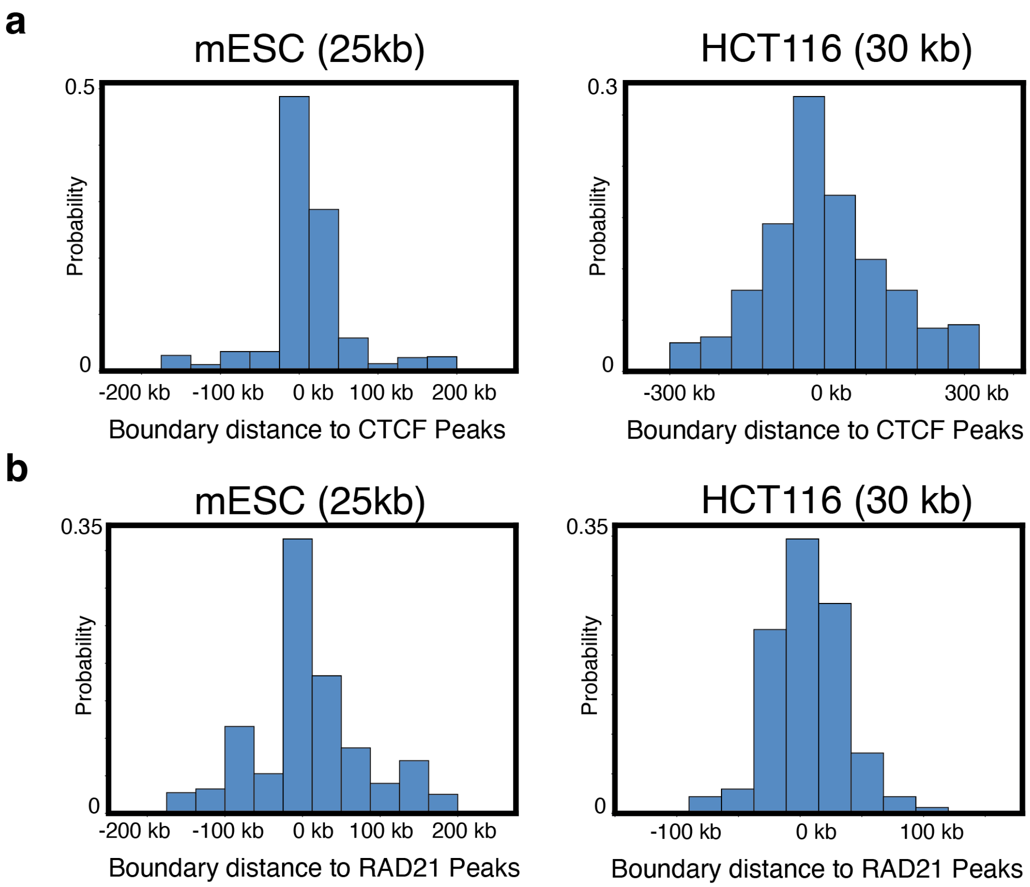


**Supplementary Figure 4. Distance of FISHnet boundary calls from CTCF and RAD21 peak calls. (a)** Histogram of the distance between FISHnet boundary calls and CTCF peak calls in mESC and HCT116 sequential oligopaints data. **(b)** Histogram of the distance between FISHnet boundary calls and RAD21 peak calls in mESC and HCT116 sequential oligopaints data.

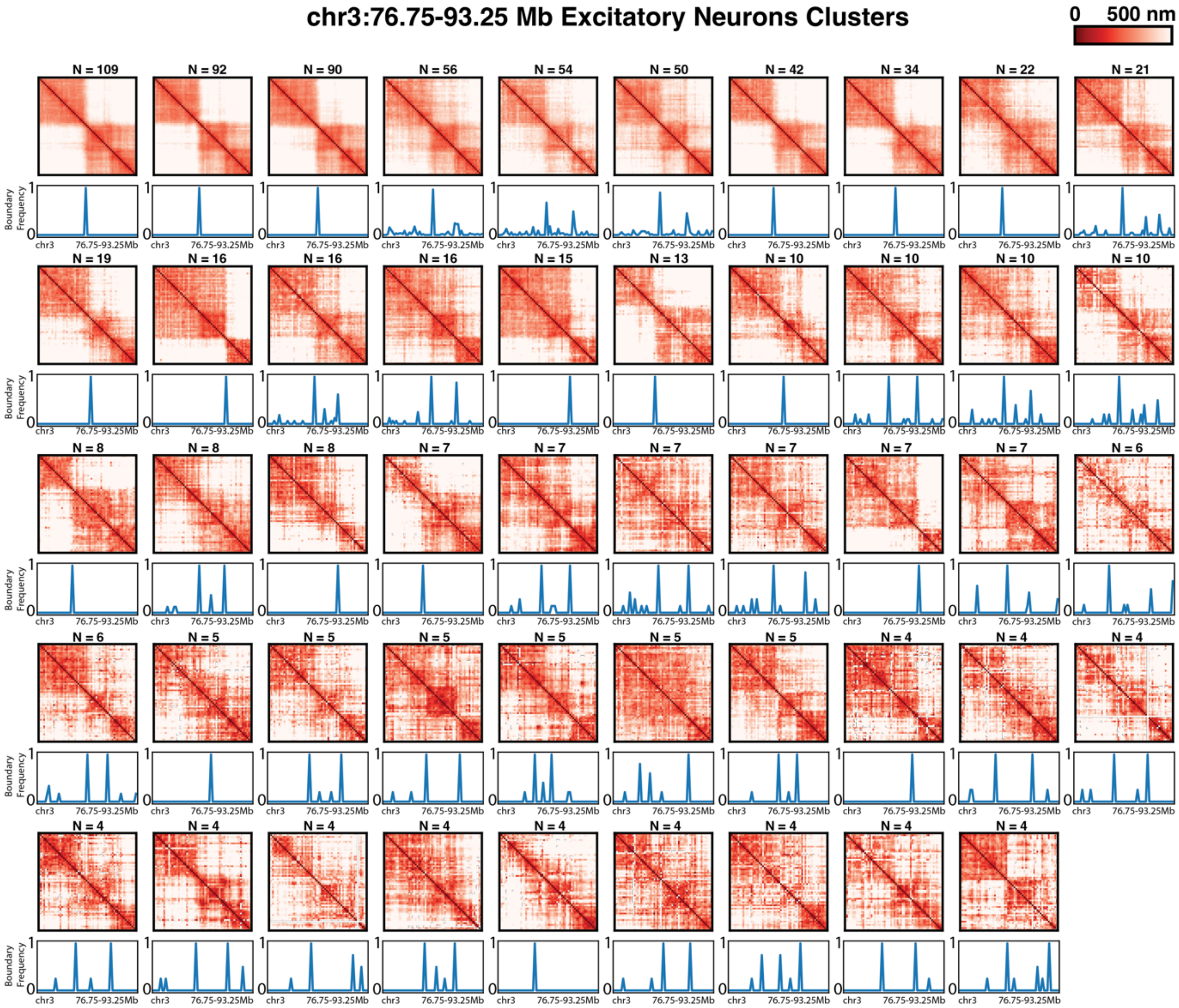


**Supplementary Figure 5. Excitatory neuron clusters created using FISHnet boundary calls.** Mean psuedo-ensemble bulk matrices for each cluster along with FISHnet frequency domain calls underlaid underneath each cluster. N indicates the number of pairwise distance matrices present in each cluster.

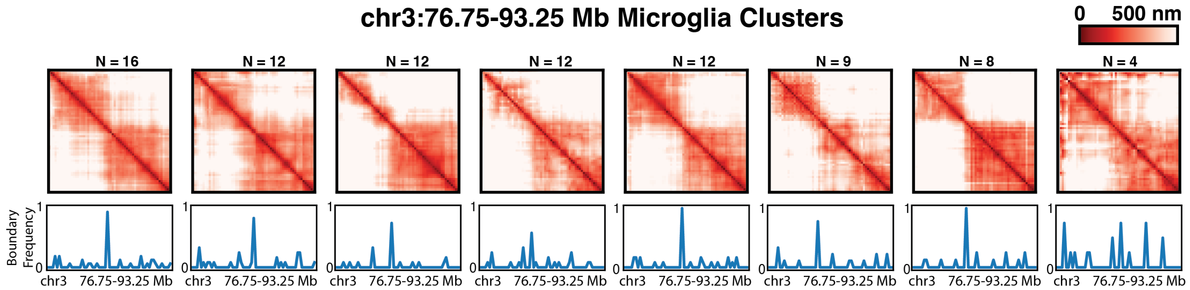


**Supplementary Figure 6. Microglia clusters created using FISHnet boundary calls.** Mean psuedo-ensemble bulk matrices for each cluster along with FISHnet frequency domain calls underlaid underneath each cluster. N indicates the number of pairwise distance matrices present in each cluster.
